# Data-driven analysis of biomedical literature suggests broad-spectrum benevolence of culinary herbs and spices

**DOI:** 10.1101/276105

**Authors:** Rakhi N K, Rudraksh Tuwani, Jagriti Mukherjee, Ganesh Bagler

## Abstract

Spices and herbs are key dietary ingredients used across cultures worldwide. Beyond their use as flavoring and coloring agents, the popularity of these aromatic plant products in culinary preparations has been attributed to their antimicrobial properties. Last few decades have witnessed an exponential growth of biomedical literature investigating the impact of spices and herbs on health, presenting an opportunity to mine for patterns from empirical evidence. Systematic investigation of empirical evidence to enumerate the health consequences of culinary herbs and spices can provide valuable insights into their therapeutic utility. We implemented a text mining protocol to assess the health impact of spices by assimilating, both, their positive and negative effects. We conclude that spices show broad-spectrum benevolence across a range of disease categories in contrast to negative effects that are comparatively narrow-spectrum. We also implement a strategy for disease-specific culinary recommendations of spices based on their therapeutic tradeoff against adverse effects. Further by integrating spice-phytochemical-disease associations, we identify bioactive spice phytochemicals potentially involved in their therapeutic effects. Our study provides a systems perspective on health effects of culinary spices and herbs with applications for dietary recommendations as well as identification of phytochemicals potentially involved in underlying molecular mechanisms.

## Author Summary

Spices and herbs are among the important ingredients in culinary preparations whose evolutionary utility is debatable. While their proximate function could be largely due to their flavor, the ultimate reason for their widespread incorporation in traditional recipes is hitherto not well understood. We implemented a computational framework for integrating spice-disease associations compiled from biomedical literature and evidence linking spice phytochemicals with diseases. By mining these tripartite data we highlight broad-spectrum therapeutic effects of spices to provide informed culinary recommendations against disease categories and seek for their potential molecular mechanisms. Through data-driven investigations, our study thus provides evidence-based applications of spices from culinary as well as medicinal perspectives.

## Introduction

Culinary practices across cultures around the world have evolved to incorporate spices and herbs in them. The potential utility of these aromatic plant products in recipes has received a lot of attention leading to multiple rationales for their wide-spread use in food preparations [1,2]. Apart from their use as flavoring agents, spices have been suggested to be of value due to their ability to inhibit or kill food-spoilage microorganisms [2]. Beyond their antimicrobial properties, the diverse therapeutic values of spices have been highlighted through *in vivo* and *in vitro* studies. Spices have been reported to possess therapeutic potential for their hypolipidemic [3], anti-diabetic [4], anti-lithogenic [5], antioxidant [6], anti-inflammatory and anticarcinogenic [7] activity.

Scientific investigations into the health effects of spices have resulted in a large body of biomedical literature mentioning their direct or indirect connections to health and diseases. With a focus on specific spice/herb, such studies have discussed their health consequences to report heterogeneous results. While some of the surveys have attempted to collate and summarize this knowledge [3,6,8], a comprehensive picture of health impacts of culinary herbs and spices based on empirical evidence still evades us. Statistics from NCBI’s PubMed data suggests an exponential increase in scientific reports associating culinary spices and herbs with diseases since 1990’s. Given their importance in food preparations, it is imperative to systematically investigate these empirical data to investigate health consequences of culinary herbs and spices.

Beyond their culinary use, traditional medicinal systems have also advocated the role of spices as therapeutic agents [8,9]. Apart from obtaining a coherent picture of the impact of these exceptional culinary ingredients on health, it would also be of value to probe the molecular mechanisms behind their action which remain largely unknown. A framework that integrates data on spice-disease associations and their phytochemicals to explore their underlying connections will help unravel molecular mechanisms behind the health impact of culinary spices and herbs (Fig 1). Towards this end, we set out to find associations between spices and diseases from biomedical abstracts available from MEDLINE using a text mining approach.

**Fig 1.**
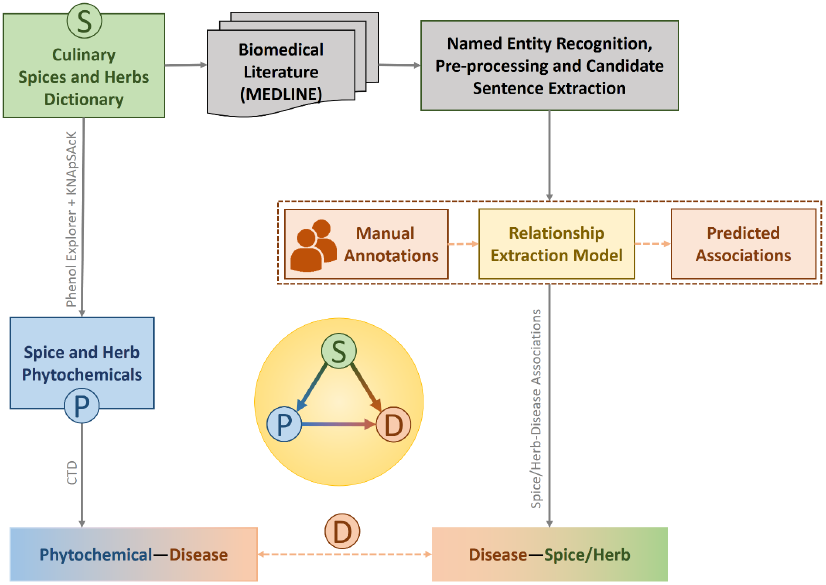
Workflow implemented for data-driven analysis of biomedical literature associating culinary spices and herbs to diseases. Starting with compilation of an exhaustive dictionary of culinary spices and herbs, towards identification of spice-disease associations, one thread of investigation involved implementation of a computational protocol for text mining of biomedical literature including named entity recognition of herbs/spices as well as diseases, pre-processing, extraction of candidate sentences, manual annotations followed by predictions of associations with a machine learning based model. The other thread involved identification of bioactive spice phytochemicals and linking them to diseases. By integrating tripartite information of spices-phytochemicals-diseases, this study establishes the broad-spectrum benevolence of spices, suggests ways for their disease-specific culinary recommendations and probes potential molecular mechanisms underlying their therapeutic properties. Thus it provides a systems perspective to health effects of spices with potential culinary and medicinal applications.

One of the earliest attempts in linking diet and diseases from literature was by Swanson who connected the utility of dietary fish oil for the treatment of Reynaud’s syndrome from indirect associations manually inferred from literature survey [10]. Biomedical literature has expanded by many folds since this pilot study making it impossible to manually concatenate the information available from research articles to infer relationships between different entities or to formulate a hypothesis. Computational approaches to text mining and natural language processing are potent tools in this pursuit [11] and many studies in recent years have contributed to efforts in this direction [12–15]. NutriChem [16] database relates plant-based foods, their phytochemicals, and diseases by using a text mining approach. HerDing [17] is another resource which links herbs to diseases by indirectly connecting constituent chemicals of the former to genes associated with the latter.

We investigated the impact of culinary spices and herbs for their role as regulators of health by text mining biomedical literature to assimilate, both, positive and negative associations. We observed that in general, the benevolent effects of spices span a broader spectrum of disorders than their adverse effects. Thus by exhaustively integrating evidence for beneficial and harmful effects of spices, we provide a framework for identification of spices whose benefits far outweigh compared to their harms. We also suggest ways for their informed culinary use as well as for identification of phytochemicals with potential therapeutic value. In summary, our study offers a systems perspective of health effects of spices and herbs to provide informed culinary recommendations and insights into underlying molecular mechanisms.

## RESULTS

### Protocol for integration of spice-phytochemical-disease data

We text mined spice-disease associations from abstracts available in MEDLINE, the largest database of biomedical literature containing more than 24 million references to research articles in biomedicine. We first manually compiled a comprehensive dictionary of 188 species of culinary spices and herbs from various sources such as FooDB (http://foodb.ca), Wikipedia (https://en.wikipedia.org/wiki/List_of_culinary_herbs_and_spices), PFAF (Plants For A Future, http://www.pfaf.org/user/Default.aspx),FPI(Food Plants International, http://foodplantsinternational.com) and FlavorDB [18] (http://cosylab.iiitd.edu.in/flavordb). This dictionary was then used to retrieve relevant abstracts from MEDLINE database. Named Entity Recognition (NER) and normalization was carried out using a dictionary matching approach for spices and NCBI’s TaggerOne [19] tool for diseases. For extracting relations, we only considered sentences that mention at least one spice/herb and disease. We manually labeled spice-disease associations for a subset of collected abstracts in order to train a machine learning classifier to predict the spice-disease associations in remaining abstracts. To further probe putative molecular mechanisms for benevolent effects of spices, we identified spice phytochemicals from PhenolExplorer [20] and KNApSAcK [21] to find their therapeutic associations with diseases using Comparative Toxicogenomic Database [22] (CTD). Fig 1 depicts the computational framework implemented for integrating and extracting tripartite spice-phytochemical-disease associations. These data are available as an integrated resource for health impacts of culinary herbs and spices [23] (http://cosylab.iiitd.edu.in/spicerx).

### Spices disease associations

We shortlisted 11750 MEDLINE abstracts containing at least one sentence with herb/spice and disease mention(s). After tagging spice/herb and disease entities, the task of relationship extraction can be treated as a multi-label classification problem with three classes of associations: positive, negative, and no associations. We used a Convolutional Neural Network (CNN) classifier with word, position, part of speech and chunk embedding [24–26] for extracting intra-sentence spice-disease relationships. It obtained an accuracy of 86.7% and macro-averaged precision, recall and F1 score of 90.7%, 80% and 84.2% respectively on an external test set. The class-wise performance metrics for the model are provided in Table 1. By combining manually annotated and predicted associations, we obtained a total of 8957 spice-disease associations from 5769 abstracts. Among these 8172 were positive spice-disease associations and 783 were negative. Out of 188 spices present in the dictionary, we obtained associations for 152 spices linking them to 848 unique disease-specific MeSH [27] (Medical Subject Headings) IDs (S1 Dataset). We observed an exponential increase in articles reporting therapeutic properties of spices after 1995 (Fig 2A).

**Fig 2.**
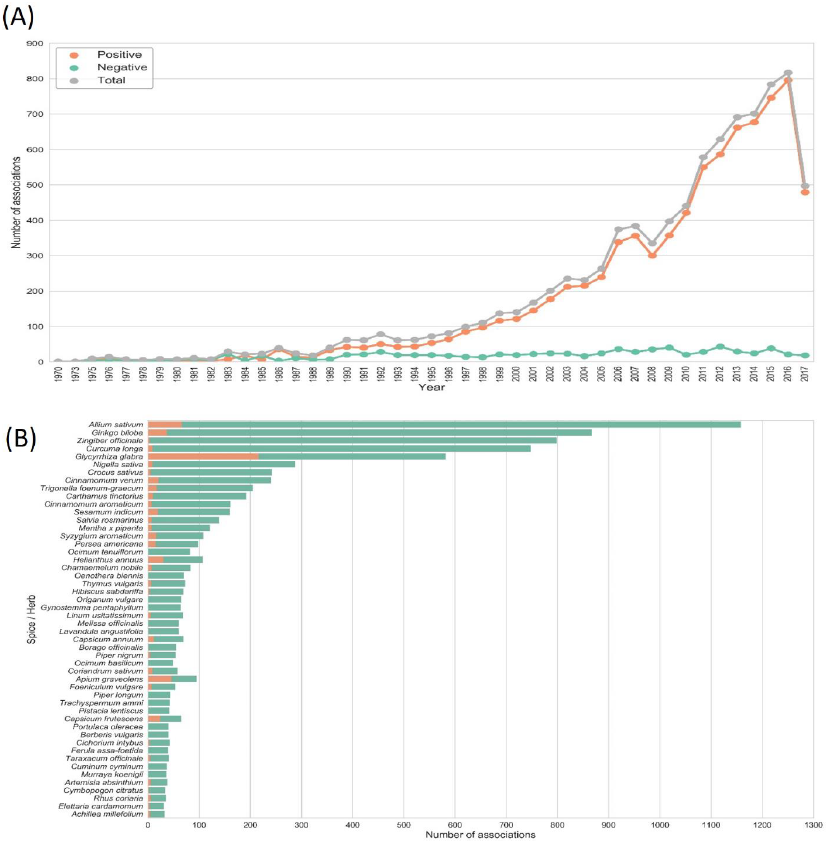
(A) Historical trend in biomedical literature reporting spice-disease associations. There is an exponential increase in articles reporting the therapeutic effects of spices in last few decades. Data of research articles archived in MEDLINE till July 2017 is represented in the illustration. **(B)** Statistics of positive and negative disease associations for the top 50 spices with most number of associations. Notice that certain spices like liquorice (Glycyrrhiza glabra) and celery (Apium graveolens) had equal number of positive as well as negative associations. The bias in number of associations may also indicate the inherent biases suggesting that certain spices are studied more than others.

**Table 1.** The class-wise performance metrics for the best CNN model, implementing word, position, part of speech and chunk embedding features, used for spice-disease relationship extraction. All negative associations were cleaned manually.

| Class | Precision | Recall | F1-Score |
| --- | --- | --- | --- |
| No-Association | 88.06% | 89.83% | 88.9% |
| Negative | 1* | 65.96% | 79.49% |
| Positive | 83.98% | 84.32% | 84.15% |
| Macro averaged | 90.68% | 80.03% | 84.20% |

\*cleaned manually

Disease entities were recognized and normalized to their corresponding MeSH IDs using TaggerOne [19]. MeSH [27] is a controlled vocabulary of biomedical terms curated and developed by National Library of Medicine. It organizes terms hierarchically from general to more specific (S1 Fig). In this hierarchical structure, a spice may have associations with a disease at multiple levels of specificity. For example, Endocrine System Diseases (C19) present at the first level of MeSH hierarchy constitutes disease sub-categories such as Adrenal Gland Diseases (C19.053), Diabetes Mellitus (C19.246) at the second level. Further, specific types of Diabetes Mellitus such as ‘Diabetes Mellitus, Type 1 (C19.246.267)’, ‘Diabetes Mellitus, Type 2 (C19.246.300)’ appear at the third level. To conduct a multi-level analysis, we associated spices with disease terms at three levels of MeSH hierarchy labeled as ‘category’, ‘sub-category’ and a ‘disease’.

We found that the number of abstracts reporting positive associations of spices with diseases far out-number those reporting negative associations (Fig 2B). A large number of spices such as ginger (*Zingiber officinale*) and turmeric (*Curcuma longa*) have very few negative associations reported in MEDLINE whereas a few others like liquorice (*Glycyrrhiza glabra*) and celery (*Apium graveolens*), have almost an equal number of abstracts reporting positive and negative associations. The complete list of associations for spices is provided in S2 Dataset. These data suggest that, in general, beneficial effects of spices have been reported more widely than their adverse effects in biomedical literature.

On analyzing individual diseases (third level of MeSH hierarchy) associated with spices, we found that diabetes mellitus, inflammation, and carcinogenesis have the highest number of positive associations (Fig 3A) (S3 Dataset). Spices were also shown to have a preventive role in various cancers including breast, colorectal, prostatic and liver neoplasms. Among the diseases adversely affected by spices were hypersensitivity, dermatitis, rhinitis, hypertension and allergic rhinitis, (Fig 3B) (S3 Dataset). It is worth noting that majority of these diseases are autoimmune in nature and are subjective to certain individuals sensitive to that spice. In such cases, spices may act as triggering factors rather than causal agents.

**Fig 3.**
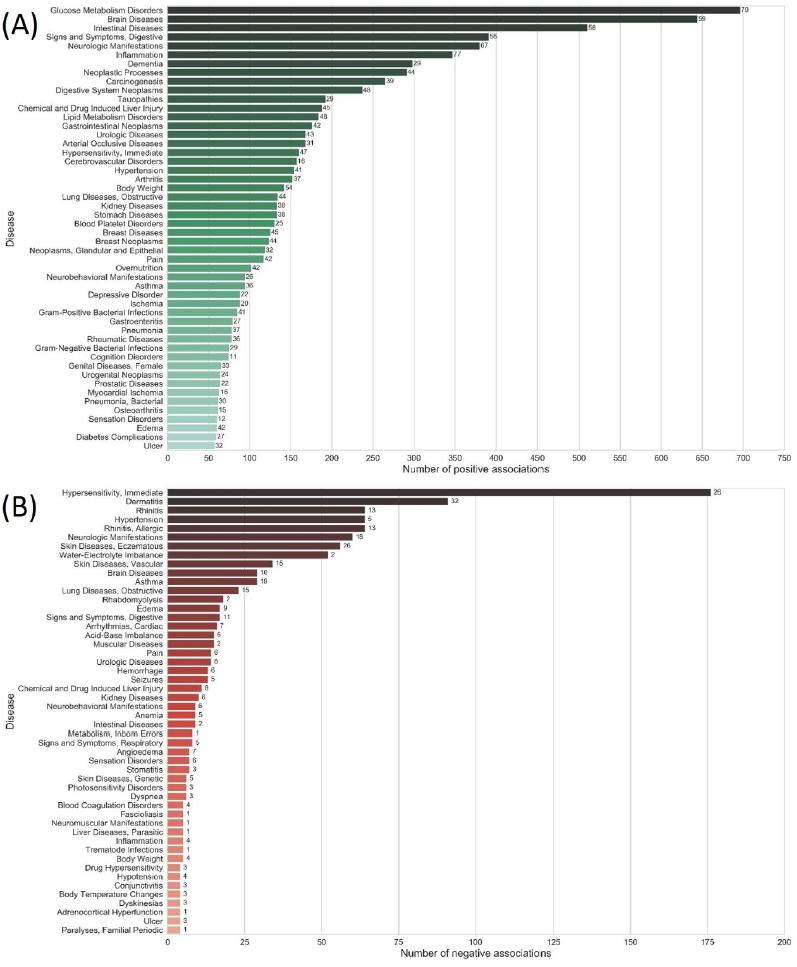
Top diseases (Third level of MeSH hierarchy) ranked according to their total number of(A) positive and (B) negative associations. Numbers shown against the bars indicate the ‘number of spices’ involved in the associations. The number of positive disease associations for spices outnumber the number of negative associations indicating that spices, in general, have been reported with beneficial health effects.

### Broad-spectrum benevolence of herbs/spices

To probe for the effects of spice/herb across a spectrum of disorders, we analyzed its associations with disease ‘sub-categories’ at the second level of MeSH hierarchy (S1 Fig). Analyzing associations at this level provides a balance between specificity and generality of disease terms. Among the disease sub-categories positively associated with spices, pathologic processes, signs and symptoms, metabolic diseases, diabetes mellitus, vascular diseases as well as central nervous system diseases were found to be dominant (Fig 4A), S4 Dataset). Top disease categories which were negatively associated with spices included vascular diseases, skin diseases, hypersensitivity and respiratory hypersensitivity (Fig 4B), S4 Dataset).

**Fig 4.**
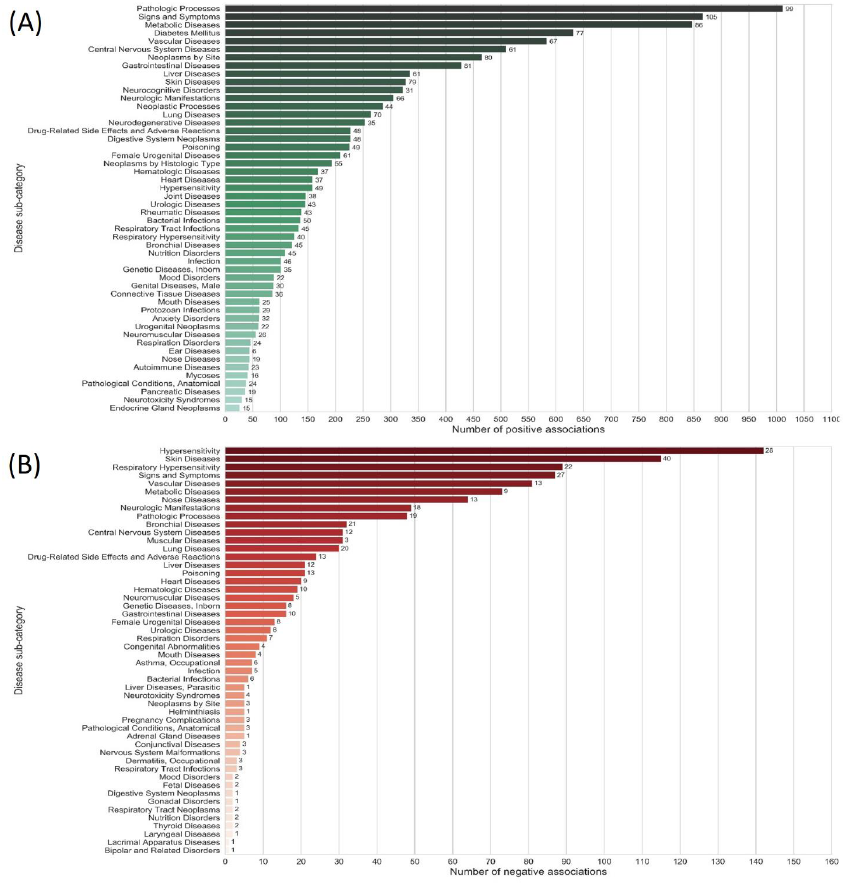
Disease categories (First level of MeSH hierarchy) ranked according to the number of (A) positive and (B) negative associations with spices. Numbers shown against the bars indicate the ‘number of spices’ the associations. The number of positive disease category associations for spices outnumber those with negative associations further confirming the benevolent health effects of spices.

To quantify the broad impact a spice may have across diverse disease categories as well as sub-categories, we devised a ‘spectrum score (*Ω*_*s*_)’ This metric computes the sum of proportion of disease terms associated at the second level of MeSH hierarchy (sub-categories), multiplied by the number of disease terms associated at the first level (categories). (See Materials and Methods). With 27 disease categories, the lower and upper bound for the spectrum score is 0 and 729 respectively. To elucidate further, let us consider a spice that is associated with all diseases in exactly half of the MeSH disease categories versus another spice that has associations with half of the diseases in every disease category. In such a case, the latter would have a higher spectrum score than the former. We computed the spectrum score for both positive (benevolence spectrum score,*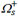* as well as negative associations (adverse spectrum score, *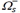*

The spices with highest ‘benevolence spectrum score’ according to our analysis were garlic (*Allium sativum*), ginger (*Zingiber officinale*), turmeric (*Curcuma longa*), liquorice (*Glycyrrhiza glabra*), ginkgo (*Ginkgo biloba*), black cumin (*Nigella sativa*), cinnamon (*Cinnamomum verum*) and saffron (*Crocus sativus*) whereas the top adverse spectrum spices were liquorice *(Glycyrrhiza glabra)*, ginger *(Zingiber officinale)*, fenugreek *(Trigonella foenum-graecum)*, ginkgo *(Ginkgo biloba)*, sunflower (*Helianthus annuus*) and *Celery (Apium graveolens).* Spices such as garlic, liquorice and ginkgo have a high benevolence as well as adverse spectrum score.

We found that for 150 out of 152 spices, the ‘benevolence spectrum score’ exceeded the ‘adverse spectrum score’, with almost 50 spices having ‘relative benevolence’ (*Δ03A9;*_*s*_) greater than 50 (Fig 5). Hence, it may be concluded that in general spices have positive effects with a broad spectrum of diseases in contrast to their negative effects which are comparatively narrow-spectrum. In line with our analysis, spices have been reported to be effective against a range of disorders [3–5,7]. Details of benevolent, adverse as well as relative benevolence scores for all spices are provided in the S5 Dataset.

**Fig 5.**
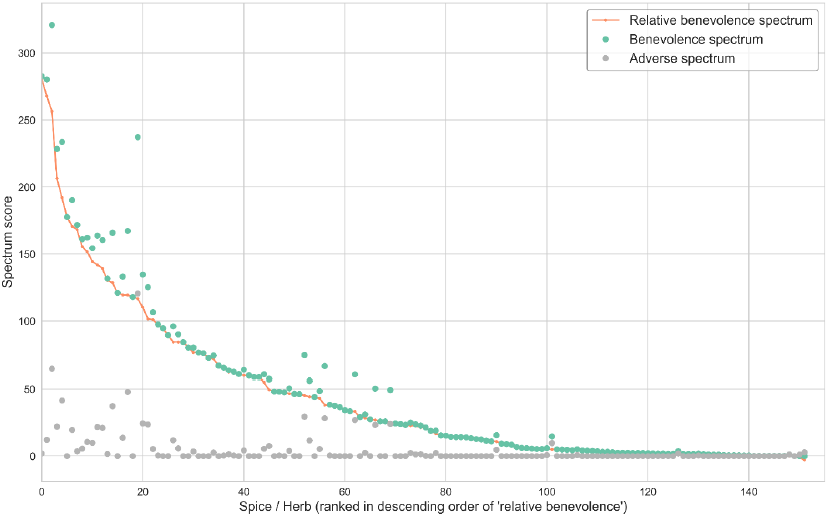
Spices ranked according to their ‘relative benevolence score’ highlighting their broad-spectrum benevolence. This score enumerates the relative health benefits as reflected in the difference between ‘benevolence spectrum’ and ‘adverse broad’ scores. Barring two, all spices had positive scores with a large number of them showing significantly larger therapeutic effects compared to their adverse effects.

### Culinary recommendations

Each of the MeSH disease categories refers to a class of disorders such as nutritional and metabolic disorders, cardiovascular diseases, nervous systems diseases, digestive system diseases, immune system diseases, neoplasms, bacterial infections and mycoses, virus diseases and such. The spectrum score forms the basis to prioritize spices for culinary intervention against a MeSH disease category. We computed the category-specific ‘benevolence spectrum’ and ‘adverse spectrum’ scores to enumerate the ‘trade-off score’ that represents the therapeutic value of a spice against a class of disorders. S6 Dataset provides a list of culinary recommendations intended as a dietary intervention against various disease categories.

There is ample amount of empirical evidence for the recommendations provided by our study. Our data suggest that spices show therapeutic effects against most of the viral diseases. Among them, turmeric (*Curcuma longa*) is the most broad-spectrum antiviral spice and is reported with inhibitory properties against various viruses including HIV, influenza, and coxsackievirus [28]. Studies in human and animal models have shown that dietary spices significantly stimulate the activities of digestive enzymes of the pancreas and small intestines such as pancreatic lipase, amylase and proteases thereby acting as digestive stimulants. Spices like ginger and garlic stimulate TRPV1, a sensor in the digestive system which has implications for gastrointestinal tract pathology and physiology [29,30]. Prominent spices recommended for cardiovascular diseases, such as tulsi (*Ocimum tenuiflorum*), mint (*Mentha X piperita*), ginkgo (*Ginkgo biloba*) and ginger (*Zingiber officinale*), have been reported with beneficial effects against cardiovascular disorders. Epidemiological studies suggest that these spices lower cholesterol level, decrease platelet aggregation, reduce blood pressure, and increases antioxidant status which in turn decreases the chances and progression of cardiovascular diseases [31]. Black cumin (*Nigella sativa*), turmeric (*Curcuma longa*) and garlic (*Allium sativum*) are the prominent spices recommended for diabetes, a major metabolic disorder. Evidence from animal studies and human trials have indicated that these spices modulate hyperglycemia and lipid profile function. Their antioxidant characteristics and effects on insulin secretion, glucose absorption, and gluconeogenesis make them potent candidates towards treating diabetes [32,33]. Similarly, the anti-diabetic property of ginkgo (*Ginkgo Biloba*) may be linked to the ability of its extract to reduce insulin resistance.

Incidentally, the spices that frequent in the culinary recommendations are among those used for culinary and medicinal preparations across cultures. Curcumin (*Curcuma longa*) and tulsi (*Ocimum tenufloreum*), widely used in Indian culinary and medicinal preparations, were present across recommendations made throughout the spectrum of MeSH disease categories. Similarly garlic, used in Southern European especially Italian cuisine, also appeared in culinary recommendations across all categories of diseases. Some of the other most potent spices include ginger (*Zingiber officinale*), black cumin (*Nigella sativa*) and ginkgo (*Ginkgo biloba*) (see S1 Table).

### Linking spices to diseases through phytochemicals

Our analysis suggests that beyond their utility as flavoring, coloring, and food preserving (antimicrobial [2]) agents, spices may have been incorporated in traditional culinary practices due to their beneficial health effects across a spectrum of disorders. Given that the therapeutic properties of plants are mediated by their phytochemicals [34–36] we hypothesize that the broad spectrum benevolence of spices can be attributed to the presence of bioactive phytochemicals such as polyphenols [36]. For example, curcumin, a polyphenol from turmeric is known to have a wide range of health benefits including antioxidant, anti-inflammatory, and anticancer effects [37]. Ajoene, a polyphenol compound derived from garlic, has been shown to induce apoptosis in leukemic cells [38]. Similarly, eugenol present in clove is reported to have antifungal property [39]. The antioxidant activity of black pepper has been attributed to the presence of β- caryophyllene, limonene, β-pinene, piperine and piperolein in its essential oil and oleoresins [39]. The anticancer properties of ginger are attributed to the presence of certain pungent vallinoids, gingerol, and paradol, as well as some other constituents like shogaols, zingerone, amongst others [39]. Going beyond the investigation of spice-disease associations, we linked spices to their constituent bioactive molecules and further connected them to diseases to obtain potential evidence of therapeutic associations (Fig 1).

We obtained 866 chemical compounds corresponding to 142 culinary spices in our dictionary from PhenolExplorer^15^ and KNApSAcK [21]. The phytochemicals data comprised of 866 chemical compounds for 142 culinary spices from our dictionary and consisted of 2042 spice-phytochemical associations. These data were filtered using PubChem [40] to keep only 570 bioactive phytochemicals, as they are known to react with tissues or cells. Further, we associated spice phytochemicals to diseases with the help of CTD [22], a public database of curated and inferred chemical-disease associations from the literature. CTD [22] classifies chemical-disease associations into therapeutic, inferred or marker associations. Therapeutic and marker associations are directly curated from the literature, whereas inferred relations are obtained from indirect associations. Therapeutic associations between a phytochemical and disease imply the presence of direct evidence of that phytochemical in alleviating the disease. For our further analysis, we focused only on 211 bioactive chemicals from the spices which were reported to have therapeutic associations.

We integrated the spice-disease associations obtained from our analysis with spice-phytochemical and phytochemical-disease mappings. This composite spice-phytochemical-disease space forms the basis for finding putative molecular mechanisms behind the benevolent effect of spice against disease. We probed for positive spice-disease associations which could be explained via the known therapeutic effects of phytochemicals reported in the spice. Out of 4380 positive spice–disease associations predicted by our model, 37% (1619) could be explained through evidence of phytochemical actions at the level of specific diseases (third level of MeSH) whereas 45% of them could be explained at the level of disease categories (first level of MeSH).

S7 Dataset provides an exhaustive list of spice-disease associations and shortlisted phytochemicals identified from the integration of tripartite data of diseases, spices and their phytochemicals (Fig 1). This analysis presents with potential molecular causal mechanisms responsible for the therapeutic effect of a spice against a disease which could be investigated further. To elucidate, garlic (*Allium sativum*) is a spice generally known for its anti-carcinogenic effects. We collected the available bioactive phytochemicals in garlic and probed for their possible therapeutic links. With the help of CTD [22], we found that allyl sulfide, a compound in garlic, is therapeutically associated with liver neoplasms. The prediction made by from our analysis linking the positive association of garlic with liver neoplasm (among other cancers) via allyl sulfide is also independently supported by the literature [41].

We probed the molecular basis for 1619 therapeutic spice-disease connections from 4380 positive associations predicted by our model. The remainder may serve as hypotheses for further investigations. Phytochemicals identified from this analysis could also be studied for their potential interactions with disease-specific genes/proteins. S8 Dataset provides the list of positive spice-disease associations for which no specific bioactive phytochemical could be obtained from our studies.

## Discussion

Humans are unique in having developed the ability to cook, which has been argued to be critical for the emergence of their large brains [42,43]. While cooked food must have provided with much-needed energy supply, it is intriguing that they flavor the food with nutritionally insignificant quantities of herbs and spices. Going beyond the ability of spices to act as flavoring and antimicrobial agents[2], our studies show the broad-spectrum benevolence of spices through analysis of empirical data. Recent studies have shown the potential benefits of consumption of spices such as chillies through cohort studies [44] as well as the role of specific spice phytochemicals in their health effects [45]. Interestingly, the broad-spectrum benevolence score of a spice was not positively correlated with its phytochemical repertoire (S2 Fig) suggesting that richness in the phytochemical content itself does not explain its therapeutic value.

We also point out negative health effects of spices, largely reflected in allergies, immune system, and skin-related disorders. Few of the negative effects of spices have been linked with their excessive use. For example, licorice, a beneficial herb for hypertension can cause weight loss, hypokalemia and other related adverse effects if consumed in large doses. Beyond probing the molecular basis of positive associations, it would also be of interest to identify toxic phytochemicals present in spices and assess their effect on specific diseases so as to provide an advisory against their consumption. Negative associations for spices projected by our study can serve as a basis for such investigations.

As opposed to a previous attempt in this direction [15,16] that linked all plant-based foods with diseases and phytochemicals from literature, our study focused on culinary spices and herbs. We investigated an exhaustive dictionary of 188 culinary herbs and spices with far better coverage (99 additional) than that of NutriChem [15,16]. Overall, in terms of the number of disease associations, the depth of our analysis was better than that of NutriChem [15,16] (S3 Fig) and our data comprised a larger set of associations for most spices (S4 Fig). NutriChem [15,16] used dictionary based string matching approach for named entity recognition and normalization of diseases as well as plants. In case of diseases, it is empirically shown that depending on the disease dictionary used, the string matching approach typically leads to a low precision and recall [46]. We used TaggerOne [19], a machine learning based named entity recognition tool which yields state of the art performance. Even though the performance of our relationship extraction model was evaluated on a dataset consisting of positive, negative and neutral associations in contrast to previous studies which evaluated on only positive and negative associations, our model achieves a comparative F1 score. In addition to this, we provide an accurate information of adverse effect of spices by manually correcting all predicted negative associations.

Similar to languages where words are synthesized from the same phonetic repertoire, cuisines around the world have concocted their own unique ingredient combinations, especially those made from spices [47,48]. Interestingly, many cuisines around the world such as those from the Indian subcontinent (*paanch phoron, garam masala, sambar masala* among a host of others referred to as *masala*), Ethiopia (*berbere*) and Middle East (*baharat*) to mention a few, have ended up developing unique spice combinations of their own. It remains to be critically examined whether these have been deliberately composed with an appreciation of therapeutic properties of spices and herbs, or are accidentally emerged constructs. Spices are frequently used as part of functional foods, for example, the Indian dish *rasam* is a concoction of different spices and has been reported to be hypoglycemic, anti-anemic and antipyretic [49]. *Sambar*, another predominantly spice-based recipe has been shown to work against prostrate cancer [50]. Traditional medicinal systems are also known to recommend spices as part of their prescriptions. *Trikatu* [51], a spice concoction made with black pepper, long pepper, and dried ginger has been advised to be of value against rheumatoid arthritis by Ayurveda, a classical traditional medicinal system from India. In Chinese traditional medicine, *Xiaoyao-san*, a combination of various spices, has been recommended for management of stress and depression-related disorders [52].

Cooking typically involves high-temperature processing via heating, boiling, frying and such. It could be argued [53] that heating is a simpler and more effective means of killing microbes, thereby refuting the antimicrobial hypothesis [2]. Other beneficial effects of spices (such as anti-diabetic, anti-carcinogenic and antioxidant and inflammatory), unearthed in this study, could not be argued against with this logic. Ironically, this argument raises another critical question: Whether the therapeutic properties and bioactivity of spice phytochemicals can sustain the intense heating processes typically involved in cooking [54]? Besides that, one of the ambiguous factors in appreciating the benevolence of spices is the distinction between the effectiveness of individual compounds vis-à-vis their synergistic actions. Apart from these aspects, there is ample scope for improvising the strategy for culinary recommendations as well as for identifications of molecular mechanisms involved in health impact of spices by including the data of quantity and disease-specific potency of their constituent phytochemicals. While raising a host of such critical questions related to dietary intake of herbs and spices, by investigating evidence from biomedical literature reporting health effects of culinary herbs and spices our data-driven analysis suggests their broad-spectrum benevolence.

## Materials and Methods

### Compilation of spices and herbs dictionary

We compiled a dictionary of 188 species of culinary spices and herbs. Scientific names and common names were obtained from Foodb (http://foodb.ca/) and Wikipedia (https://en.wikipedia.org/wiki/List_of_culinary_herbs_and_spices). All scientific names were mapped to their respective NCBI Taxonomy IDs, collapsing varieties wherever available, to their corresponding species ID. For example, *Capsicum baccatum var. pendulum*, the scientific name of Peruvian pepper, was mapped to *Capsicum baccatum*. This dictionary was further enriched by adding common names from FPI (Food Plants International, <http://foodplantsinternational.com/plants/), NCBI> Taxonomy (https://www.ncbi.nlm.nih.gov/taxonomy) and PFAF (Plants for a Future, http://www.pfaf.org). Singular and plural forms of common names of the spices and herbs were also included. Common names that did not exclusively map to an NCBI Taxonomy ID were ignored.

### Biomedical literature

We used MEDLINE (Medical Literature Analysis and Retrieval System Online, https://www.nlm.nih.gov/bsd/mms/medlineelements.html) as our source of biomedical literature. It includes citations from more than 5600 scholarly journals with over 24 million references to peer-reviewed biomedical and life science research articles from as early as 1946. The data was downloaded in bulk from the FTP server of NCBI (https://www.nlm.nih.gov/databases/download/pubmed_medline.html). This data is available in XML format with each file containing 30000 articles. Parsing was carried out using a modified version of PubMed parser (https://github.com/titipata/pubmed_parser) to extract information of PMID, Date, Title, Abstract, Journal, and Authors. The modified parser is available at https://github.com/cosylabiiit/pubmed_parser. Articles for which no abstract text was available were not considered.

### Named Entity Recognition

We adopted a dictionary matching approach for Named Entity Recognition (NER) of spices and herbs. With a large dictionary, the process of dictionary matching becomes a computational bottleneck. Thus to save computational time, we used a modified implementation of Aho-Corasick algorithm (NoAho, https://github.com/JDonner/NoAho) for obtaining non-overlapping and longest matches at the token level. For disease NER (DNER) and normalization, we used TaggerOne [19] which utilizes semi-Markov models with a rich feature set. It was reported to have a precision of 85% and a recall of 80% on the Biocreative V Chemical Disease Relation test set [46]. We used the pre-trained disease-only model available with TaggerOne [19] on our data.

### Preprocessing

Sentence segmentation was carried out on the retrieved abstracts using Stanford CoreNLP package [55]. Only sentences with mention of at least one herb/spice and one disease were considered for extracting relations. Complex sentences with multiple mentions of herbs/spices and/or diseases were simplified by creating multiple copies of the same sentence, and iteratively tagging a given pair of herb/spice and disease only. In all sentences, numbers were replaced by a standard identifier token and, barring some punctuation characters (!,.:;), all special characters were removed. The preprocessed sentences were then tokenized using GENIA [56] and the part-of-speech (PoS) tag, as well as the chunk tag of each token were obtained. We also computed the distance of each token from the tagged entities and used them as position features for the machine learning model.

### Labelling associations

Manual sentence-level annotations were performed for a random set of abstracts, which was used for the training as well as for evaluation of the relationship extraction model. Hitherto, to the best of our knowledge, no gold standard corpus exists for plant-based foods and diseases. We manually annotated a total of 6712 sentences to tag positive, negative and neutral spice-disease associations. Out of the total annotated data, 2669 sentences had positive whereas 301 had negative associations. No associations were found in the rest of the sentences.

### Machine learning based computational models

Starting with preprocessed and manually labeled sentences, we developed a machine learning classifier for extracting relations from unlabeled sentences. We tested the following models[24–26]: (i) Linear Support Vector Machine (SVM) with unigram and bigram word features; (ii) Convolutional Neural Network (CNN) with word embedding[26] features; and (iii) CNN with word, position, PoS and chunk embedding.

For the Linear SVM model, we obtained the unigram and bigram word features and scaled their respective weights using Term Frequency-Inverse Document Frequency (TF-IDF) approach. This model was trained using one-versus-all strategy. Following are the equations describing the method for computing TF-IDF weights of features: (*i*)*tf* (*t,s*) = *f* _t,s;_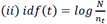;and (*iii*) *tfidf* (*t,s*) =*tf (t,s)* \**idf(t)* where, *f*_*t,s*_denotes the number of times feature *t* appears in sentence *s,n*_*t*_ is the number of sentences in which the feature appears and is the total number of sentences.

For both the CNN models, the word embedding was initialized using pre-trained weights from Chiu *et. al* [57]. The embedding for unknown words was initialized from a uniform (-*α, α*) distribution, where *α*was determined such that the unknown vectors have approximately the same variance as that of pre-trained data [58]. For the second CNN model, the PoS and chunk features were obtained using GENIA[56] tagger and mapped to a randomly initialized embedding matrix. Similar mapping was performed for the position features which include the distance of each token from the spice-entity as well as the disease-entity. Further, sentences were zero-padded to equalize their lengths and different features for each token were concatenated to form a single vector representation. The first layer of both the CNNs thus transforms each sentence into a *d*×*n* matrix, where *d* is the number of dimensions of the ‘token’ vector of the sentence and *n* is the maximum sentence length in the whole corpus. The two CNN networks differ only in the value of *d* The second layer is a Convolutional layer with *n*_*f*_ filters of different filter sizes *f*_*z*_ and rectified linear unit (ReLU) activation. The respective maximum activations of all the filters are then concatenated into a single vector of size *n*_*f*_ and fed to a Dropout layer [59], which randomly sets an activation to zero with probability.*p* This is followed by a dense layer of *n*_*h*_hidden units with ReLU activation and a softmax layer with 3 units. For both the networks, we used categorical cross entropy as our objective function and applied *l*_2_ regularization of 3 on the dense layers. The networks were trained using mini-batch gradient descent with shuffled batches of size 50 and Adam [60] optimizer. We adopted an early stopping criterion for the training process and stopped model training if the validation loss did not decrease for 5 epochs. To address the class imbalance problem, we over-sampled the negative class and the positive class by a factor of 12 and 1.35 respectively. The hyper-parameters of both the neural networks were determined using 5-fold cross validation and are available in S2 Table.

### Evaluation metrics

We evaluated the performance of our model based on its precision, recall, F1 score and accuracy: *precision=TP/(TP+FP);Recall=TP/(TP+FN);F1-score=2. Precision-Recall/(precision+ Recall)Accuracy=TP+TN/(TP+TN+FP+FN)*,whereare *TP,FP,TN,FN* are True Positives, False Positives, True Negatives and False Negatives respectively.

### MeSH hierarchy

MeSH is a controlled vocabulary of biomedical terms curated and developed by National Library of Medicine. The terms are hierarchically organized from generic to more specific. The DNER tool used in this study (TaggerOne [19]) normalizes the tagged entities to MeSH IDs. The hierarchical structure of MeSH results in situations where a spice is typically associated with a disease at multiple levels of specificity. For example, in the first level of MeSH hierarchy a spice may be linked with the disease category Endocrine System Diseases (C19) and at the second level C19 may be associated with sub-categories such as Adrenal Gland Diseases (C19.053) or Diabetes Mellitus (C19.246). Further, it may be linked to the specific type of Diabetes Mellitus, say, ‘Diabetes Mellitus, Type 1 (C19.246.267)’ or ‘Diabetes Mellitus, Type 2 (C19.246.300)’ appearing at the third level. We conducted a multi-level analysis by associating spices with disease terms at top three levels of MeSH hierarchy which were referred to as category, sub-category, and a disease (S1 Fig).

### Adverse and benevolent spectrum scores

The ‘spectrum score of a spice (*Ω*_*s*_)’ encodes diversity of adverse *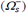* or therapeutic *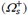* effects of a spice *s* across the MeSH disease categories as well as their constituent subcategories, and is defined as *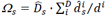.* Here, is total number of MeSH disease categories, *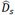* represents the number of disease categories with which spice has therapeutic association with, *d*^*i*^ represents the total number of disease sub-categories in th *i* th disease category, and *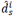* represents the number of disease subcategories in the th disease category with which the spice *s* is associated. When calculating the ‘spectrum scores’ across all 27 categories, the ‘adverse spectrum score’ and ‘benevolent spectrum score’ vary between 0 and 729. Further, for each spice the ‘relative benevolence’*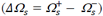* that encodes its residual therapeutic benefit was computed.

**‘Therapeutic tradeoff score’ for culinary recommendations**

Category-specific (benevolence and adverse) spectrum score was defined as *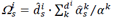.* Here, *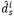* represents the number of disease sub-categories in the th disease category with which spice has therapeutic association with,*α*^*k*^ represents the total number of diseases in the *k*th disease sub-category, and *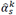* represents the number of disease subcategories in the *k*th disease sub-category with which the spice is associated. The ‘therapeutic tradeoff score’, *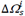*, represents the difference between the ‘benevolence spectrum’ and ‘adverse spectrum’ of spice for category *i*; the higher the tradeoff score of a spice the better is its therapeutic value against the spectrum of diseases represented by this category. Thus, tradeoff score of a spice serves as a basis for its recommendation against a MeSH disease category.

### Linking phytochemicals from spices/herbs to diseases

We obtained the phytochemical information for spices/herbs using KNApSAcK [21] and CTD [22]. The different compound identifiers were standardized to PubChem IDs and further PubChem BioAssay [40] was used for ascertaining their bioactive status. Therapeutic associations of a compound were obtained from CTD [22] after mapping its PubChem ID to corresponding MeSH ID.

## Acknowledgement

G.B. thanks the Indraprastha Institute of Information Technology (IIIT-Delhi) for providing computational facilities and support. R.N.K. thanks the Ministry of Human Resource Development, Government of India and Indian Institute of Technology Jodhpur for the senior research fellowship. R.T. (Research Associate) and J.M. (Research Intern) are affiliated to Dr. Bagler’s lab at the Center for Computational Biology, and are thankful to IIIT-Delhi for the support.

## Author Contributions

G.B. conceived the idea and supervised the work; G.B., R.T. and R.N.K. designed and performed the experiments; R.N.K and R.T. performed the data collection and cleaning; R.T. implemented the computational models; G.B., R.T. and R.N.K analyzed the results and wrote the manuscript; R.N.K. and J.M. performed the annotations; All the authors reviewed and approved the final manuscript.

## Supporting information captions

**S1 Fig: Hierarchical Structure of MeSH disease headers**. For the purpose of multi-level analysis, spices were associated with disease terms at three levels of MeSH hierarchy— ‘category’, ‘sub-category’ and a ‘disease’.

**S2 Fig: Correlation between the number of phytochemicals in spices and their broad-spectrum benevolence-** The data indicate that the broad-spectrum benevolence score of spices and their phytochemical repertoire are not correlated.

**S3 Fig: Comparison of the number of associations obtained for spices reported by our study with that of NutriChem**[16,61] **indicating richer associations in our data**

**S4 Fig: Comparison of associations retrieved for ‘individual spices’ by NutriChem**[16,61] **to those from our study, suggesting better depth/coverage in the latter**

**S1 Table:** Top ten broad spectrum spices and number of MeSH disease categories and subcategories with which they are positively associated

**S2 Table:** Hyper-parameters selected for the convolutional neural network Model 2 and Model 3.

**S1 Dataset:** Statistics of positive and negative spice-disease associations for each spice

**S2 Dataset:** Statistics of positive and negative associations as well as number of spices, at the third level of MeSH

**S3 Dataset:** Statistics of positive and negative associations as well as the number of spices at the third level of MeSH disease hierarchy.

**S4 Dataset:** S4Statistics of positive and negative associations as well as the number of spices at the second level (sub-category) of MeSH disease hierarchy.

**S5 Dataset:** Benevolent, adverse as well as relative benevolence scores for all spices

**S6 Dataset**: List of culinary recommendations against various disease categories.

**S7 Dataset**: Tripartite associations for a spice and a disease along with specific phytochemicals reported to be involved in the therapeutic action.

**S8 Dataset:** Statistics of spice-disease associations for which bo specific phytochemicals were ascertained.

